# Explaining the pathogenesis of African swine fever using knowledge-driven regulatory network modeling

**DOI:** 10.64898/2026.06.26.734812

**Authors:** Joyce Reimer, Pranta Saha, Kira Comfort, Zahed Khatooni, Heather L. Wilson, Connor Burbridge, Brook Byrns, Steven Rayan, Sureesh Tikoo, Gordon Broderick

## Abstract

A lethal DNA virus with significant economic impact on livestock farmers worldwide, the pathogenesis of African swine fever virus (ASFV) infection is complex and continues to challenge the development of effective vaccine candidates. The requirement for high-containment conditions further complicates its study, resulting in limitations in sample size and marker assessment that challenge conventional statistical analysis. In this work we demonstrate how prior knowledge of immune biology and pathogen-host proteome interactions can be leveraged and reconciled with sparse experimental data to deliver plausible mechanistically informed hypotheses describing ASFV illness progression. We apply large-scale automated mining of literature and pathway schema together with generative artificial intelligence (AI) to create closed-loop regulatory network models consisting of 133 pathogen and host proteins linked by 676 regulatory interactions. Immune regulatory tuning of these networks is reverse engineered to explain two distinct experimentally observed illness progression trajectories in only 5 markers measured every second day over a maximum of 8 days. Comparison of network model pools specific to each progression phenotype suggest that these significantly different outcomes may arise from altered regulatory tuning of genes coding for interleukin (IL)1β, tumor necrosis factor (TNF)α and Forkhead box protein (FOX)O4, potentially as a result of epigenetic adaptations. Simulated challenges with individual ASFV protein confirm broadly delayed interferon (IFN)-γI response in both phenotypes, with multigene family (MGF)505-3R offering the earliest induction and only in the more severe phenotype. Paradoxically, predictions suggest that this delay is preceded by an early IL-10 induction by this same viral protein. While added model granularity and validation is needed, we propose that this proof-of-concept knowledge driven approach offers an attractive solution to mechanistic hypothesis generation in data poor environments.

## Introduction

African swine fever (ASF) poses a significant threat to domestic and wild pig populations worldwide. First reported in 1921, it began to spread outside of Africa in 2007, and since the World Organisation for Animal Health (WOAH) began monitoring the situation in 2005, cases have been reported in 83 countries [1]. Characterized by acute hemorrhagic fever and near-100% mortality rates, the deadly illness continues to have detrimental effects on local pig farms and regional ecosystem health, as well as on global trade and food security at an international scale [1-3]. The pathogen responsible for the lethal infection is African Swine Fever virus (ASFV), a large, double-stranded DNA virus of *Asfarviridae* family that is complex and consequently difficult to characterize [4]. Because of this, the task of developing a vaccine has been effortful, yet, so far, without reward as there are currently no ASFV vaccines that are internationally recognized as safe and effective. Some live attenuated vaccines have shown host protection; however, these tend to be genetically unstable, offer very limited immunity, or they are accompanied by severe side effects [5-9]. Protein subunit vaccines are considered a viable alternative but come with their own challenges due to a lack of knowledge about the nature of the numerous possible interactions between viral and host proteomes [7]. Certainly, there has been significant research effort in the field, but the complexity of the problem is such that integrating these numerous disparate studies and distilling the latter into a collective and overarching understanding that can inform on actionable steps forward remains a challenge. Integrative knowledge engineering strategies have evolved dramatically and have shown promise in delivering mechanistically informed models in environments where data might be sparse, distributed across many studies, and where the number of observable markers is limited. This relatively low level of data is especially common in experimental environments operating under the high containment conditions required to study dangerous pathogens such as ASFV. Indeed, digital twins constructed from qualitative descriptions of causal mechanisms extracted from our growing knowledge of host biology have shown promise in explaining the event-driven dynamics of onset and progression in complex illness [10]. Large-scale semantic text-mining of peer-reviewed literature [11] and extraction for manually curated databases [12] have evolved considerably and together with the careful adoption of emerging technologies like generative AI [13,14] these technologies are driving a growing development and use of digital-twin models in healthcare and biomedical research [15,16]. These advances in modeling host biology coincide with the increasing availability of pathogen-host protein-protein interaction networks, generated with high-throughput genetic screening and data analysis [17-19]. Together these maps of pathogen-host and host immune interactions offer a foundation for describing the onset and progression of infection in terms of basic molecular mechanisms of immune response.

In this work, we apply this knowledge-driven strategy to describe the onset and diverging evolution of infection with ASFV in two otherwise identical pigs using sparse sampling of a very limited set of markers. Specifically, we use an existing ASFV-host protein-protein interaction map generated from statistical analysis of amino acid sequence alignment [17] combined with a mechanistically informed regulatory network of host immune signaling assembled with the automated mining of the literature and curated databases. Gaps in our current knowledge are addressed using generative AI under strict reliability criteria to produce a closed-loop pathogen-host regulatory network. A large-scale global optimization strategy [20] is then applied to identify competing sets of immune kinetic and decisional logic parameters capable supporting network dynamic behaviors that include the observed illness courses of interest, namely the progression of 5 distinct cytokine markers measured daily for 6 days and 8 days post-infection in two contrasting severity phenotypes [21]. Based on the hypothesis that these differences in progression might result from epigenetic adaptations, two corresponding families of models were derived and analyzed. In comparing the regulatory kinetic parameters in each phenotype’s pool of models, we find the regulation of 3 immune mediators in particular stand out as determining factors in the course of illness. Specifically, we find that blunted regulatory response in genes coding for interleukin (IL)-1β, tumor necrosis factor alpha (TNFα) and Forkhead box protein (FOX)O4 support slower illness progression in addition to other regulatory alterations. Though not definitive, we propose that the assembly and comparison of phenotype-specific logic models may provide valuable insight into potentially key epigenetic alterations to regulatory axes and mediators and that such alterations may confer increased vulnerability or resistance to illness progression. Importantly such changes in the regulatory fabric may not be otherwise visible using a more conventional statistical analysis especially given limited experimental data collected in small study populations such as this one. Such insight can be leveraged to significantly shorten the discovery cycle, shedding light on ASF pathogenesis, highlighting potential targets for ASF vaccines, demonstrating the value of examining experimental data through the lens of our prior knowledge of host biology.

## Results

### From interactome to regulatory network

The ASFV pathogen-host network used in this study was derived from the protein-protein interactome published by Wu et al. [17] which consisted initially of 77 viral proteins and 590 host proteins, for a total of 667 protein nodes linked by 8946 undirected interactions. The expected degree of protein-protein interaction affinity was estimated using an alignment frequency score, with higher scores suggesting higher affinity. Applying a thresholding technique to this frequency-based affinity score that is commonly used to filter background from foreground in image analysis, this initial protein node count was reduced to 10 viral proteins interacting with 25 host proteins linked by 34 undirected pathogen-host interactions bearing a frequency score of 540,663 or greater. Undirected relationships were translated into pairs of opposing directed regulatory relationships with an unknown mode of action to be derived through reconciliation with available data (e.g., “Protein A regulates Protein B” rather than “Protein A and Protein B are associated”). Regulatory relationships linking these 25 gateway host proteins as well as their intermediate regulators were then extracted from manually curated pathway databases to provide a basic prior knowledge structure. To support dynamic stability the final network must form a closed loop regulatory system. To achieve this closed-loop architecture, several iterations of pruning and augmentation were conducted by combining extraction from manually curated databases and with generative AI predictions of putative undocumented relationships as outlined in detail in the Methods section. This iterative process converged to produce a closed-loop, directed regulatory network consisting of 123 host proteins and 10 viral proteins, linked by 676 directed regulatory relationships (**Fig. 1**). Of these relationships, 357 upregulated the downstream target and 222 downregulated the downstream target while the mode of action for the remaining 97 relationships could not be readily extracted from documentation or predicted with confidence. With an overall connection density of ∼4%, relationships were distributed such that tightly connected areas supported a clustering coefficient of 0.14. Component nodes in this network were separated on average by no more than 3 intermediate regulators (characteristic path length ∼4 path segments), with the most distal regulators spanning 10 cascading regulatory relationships (network diameter). With a complement of 9 direct neighbors, a typical protein node in this network is mediated on average by 5 upstream regulators (median 4), with the activation of IL1β being directed by as many as 45 upstream regulators (27 upregulating, 14 downregulating, 4 unknown mode of action), followed by PTGS2 and TNFα with 23 and 20 upstream regulators respectively. This network served as the basic architecture in the subsequent identification of parameter values that dictate the regulatory decisional logic at each node and the scheduling of state transition events with the resulting data-aligned network variants serving to support the analysis of predicted phenotypic variation.

**Figure. 1:**
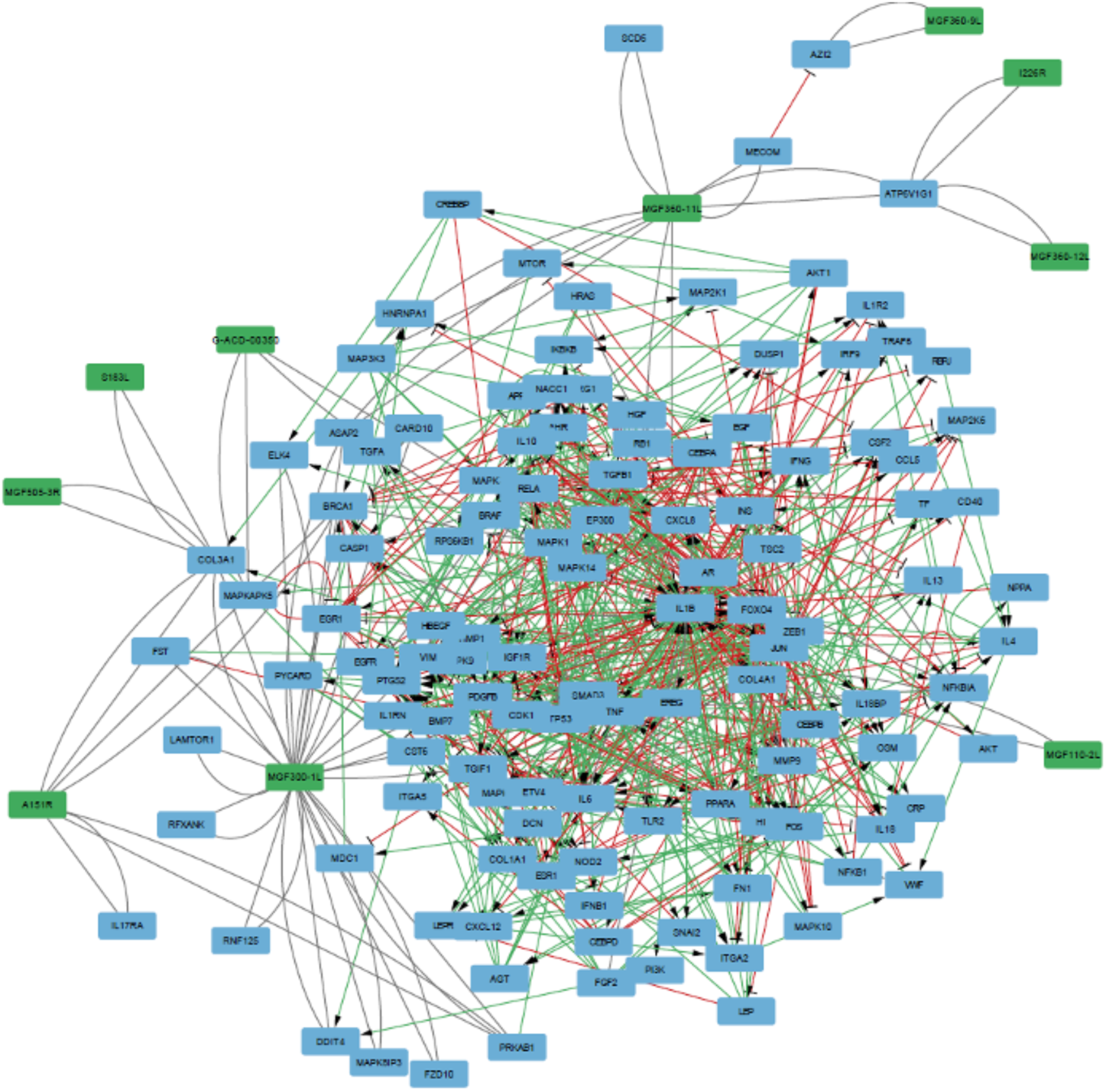
Structure of the closed-loop host-pathogen ASFV network. The network is made up of 133 nodes and 676 relations. A green node indicates a viral protein (there are 10), blue nodes are host proteins. Green lines indicate a positive regulatory relationship, red indicates inhibitory, and gray indicates unknown polarity.

**Table 1:**
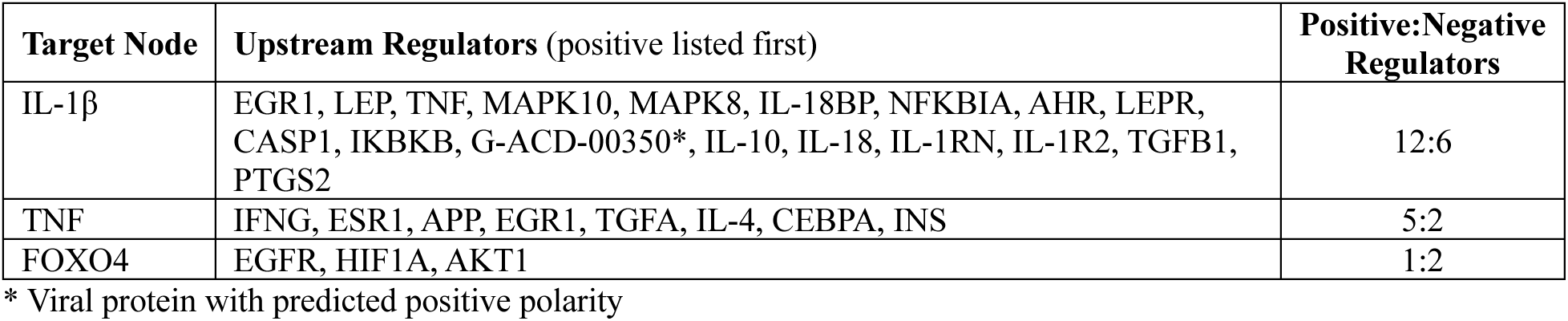
Differentially regulated relations (Slow-protective score <-20). Ratios of positive to negative regulation of target nodes that have a slow-protective score <-20. For IL-1β and TNFα, two-thirds of slowed regulators are positive; for FOXO4, two-thirds are slowed negative regulators.

| Target Node | Upstream Regulators (positive listed first) | Positive:Negative Regulators |
| --- | --- | --- |
| IL-1β | EGR1, LEP, TNF, MAPK10, MAPK8, IL-18BP, NFKBIA, AHR, LEPR, CASP1, IKBKB, G-ACD-00350*, IL-10, IL-18, IL-1RN, IL-1R2, TGFB1, PTGS2 | 12:6 |
| TNF | IFNG, ESR1, APP, EGR1, TGFA, IL-4, CEBPA, INS | 5:2 |
| FOXO4 | EGFR, HIF1A, AKT1 | 1:2 |

\* Viral protein with predicted positive polarity

### Identifying regulatory programming

With a regulatory wiring in place in the form of a foundational prior-knowledge network structure, a regulatory program must now be identified that directs flow of regulatory information through the network, namely a decisional logic and scheduling that governs the activation and inactivation of protein nodes as well as the timing of the corresponding regulatory events. Values for the detection thresholds and contextual weights of upstream regulators, the mode of action of such regulators when undocumented, as well as the allowable incremental gain in activation, the decisional lag, the rate of change (speed) and the states of unobserved protein nodes (i.e. missing data) are estimated by formulating and solving a combinatorial optimization problem such that valid parameter sets support predicted network behaviors that include what little experimental data might be available. As described in the Methods section, this parameter value search is conducted using a variant of simulated annealing [20]. For this network, the total number of parameters that comprise a model is 2,427, however, because there are 579 of 676 relations for which polarity was already known, it was only necessary to search for optimal settings in 1,848 parameters. Given the structural and data-imposed bounds defined in the Methods section, this gives rise to a search space of 9 x 10^835^ possible combinations.

In the current work, we define a valid model as one capable of adequately reproducing the observed trajectories of a subset of cytokines in swine peripheral blood following infection with ASFV. Specifically, we use the trajectories of IL-1β, IL-4, IL-6, IL-10, and TNFα measured in peripheral blood serum and the overall viral nucleic acid volumes reported by Zuo et al. [21] as a reference against which to measure model adequacy. The study distinguishes between two different cytokine trajectories, one supporting a survival of 8 days and the other a survival of 6 days. We propose that these trajectories may arise from acquired epigenetic changes defining “Less Severe” (LSev) and “More Severe” (MSev) phenotypes respectively, and consequently we identify two separate phenotype-specific model pools. Optimal sets of model parameter values are identified that minimize penalty and maximize reward, which is subtracted from the penalty to produce a net fitness. In this case, penalties are levied based on the degree of departure from the observed reference trajectories (penalty) while the reward consists of a soft constraint whereby the averaged viral protein trajectories follow a growth curve (see Methods for formulation of the objective function). Upon convergence, these optimal parameter value sets supported a minimum objective function score of -27 with a departure of only 1 bit (from a possible 30-bit departure or ∼3%) in the case of the LSev phenotype. In the case of the MSev phenotype, parameter value sets were found that supported an objective function score of -28 with a departure of 2 bits (from a possible 26-bit departure or ∼8%). The corresponding predicted cytokine trajectories traces for each phenotype are shown in **Fig. 2**, alongside the reference experimental data. Despite a non-zero departure, the simulated data closely follows the reported experimental trajectories, and many of the viral proteins also adhere to a smooth growth curve.

**Figure 2.**
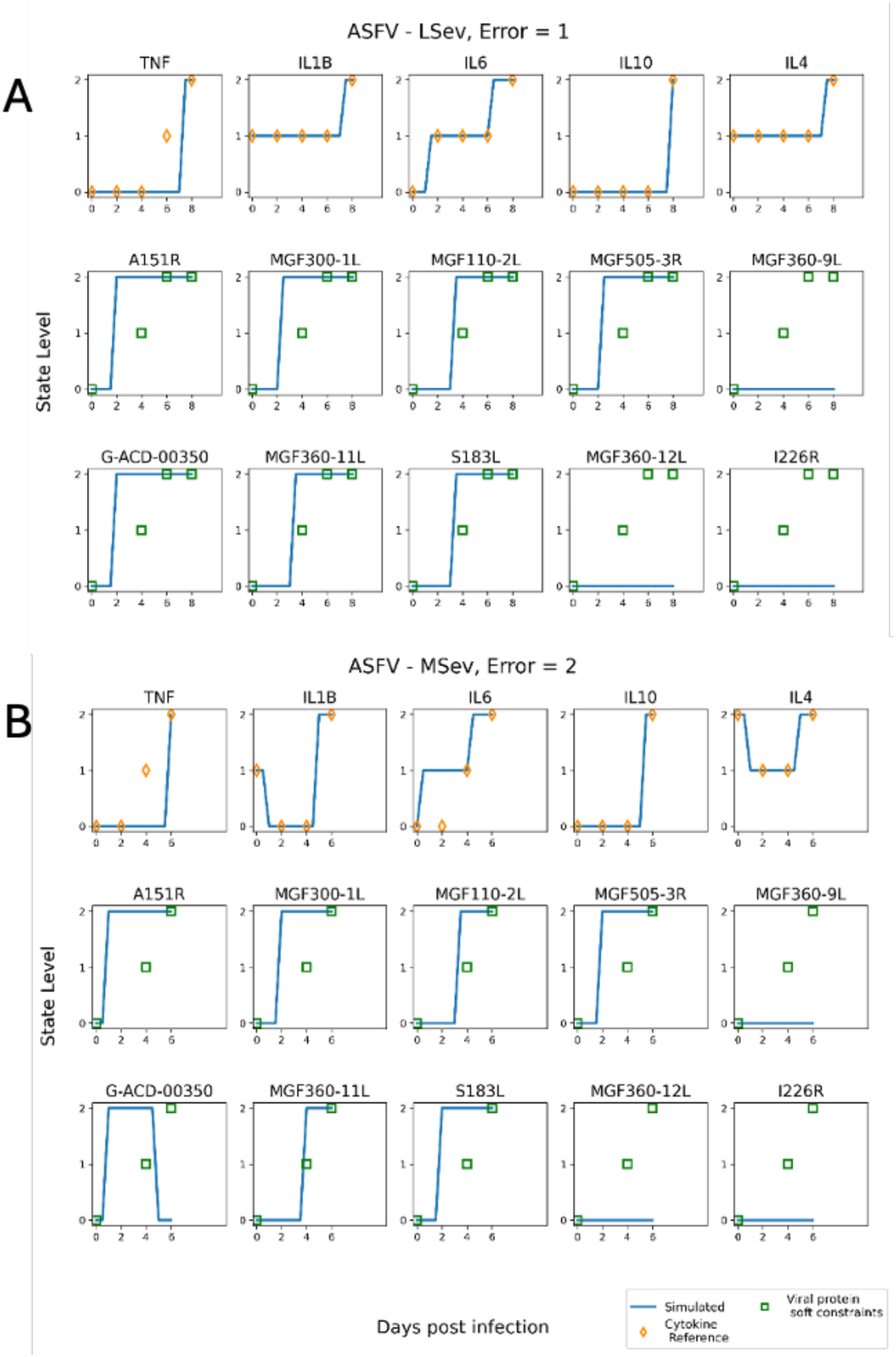
Simulated traces from lowest-error models in comparison with experimental data. (**A**) Parameter optimization for the LSev phenotype model resulted in a solution with error of 1. The traces closely match the experimental cytokine reference markers. The majority of viral proteins adhere to the soft constraints, exhibiting a typical growth curve. (**B**) The MSev phenotype model resulted in an error of 2, capturing the experimental data fairly well. The average viral protein trajectory is a typical exponential growth curve.

### Comparing phenotype-specific immune regulatory networks

With such a large search space (∼10^836^ combinations) and such a sparse reference dataset, namely 5 modeled markers observed at 4 and 5 time points in the MSev and LSev phenotypes respectively, we can expect a relatively large basin of equivalent high fitness low-departure solutions. Therefore, upon convergence, the first 100,000 distinct solutions for each phenotype that supported a departure and objective function value equal to the best minimum solution were collected and analyzed for statistical differences. Interestingly, despite this large number of competing models, the latter share strong commonalities in structure and regulatory programming within each phenotype. Specifically, in the pool of models supporting an LSev illness progression, 25% of parameters are set to identical values across all models, and 91% of all parameters are set to the same values in ≥98% of the model pool. Likewise, for the MSev phenotype 26% of all parameters are unanimously set to identical values, with 97% of parameter settings being shared across ≥98% of the model pool. These data would suggest a broad agreement in regulatory domain across models with a relatively small pool of complementary behaviors specific to unindividual models. In other words, universal agreement on at least ∼25% of the regulatory programming is required to support the same common experimental outcome observed in each phenotype, at least as it applies to the trajectory of the 5 immune markers reported.

#### Altered tuning of host immune circuitry

In distinguishing between phenotypes, we focused first and foremost on parameters related to the detection threshold for each regulatory action (edge) and the speed with which the resultant state of each component protein node is updated. Together these parameters govern the immediacy with which each protein node changes state, and we propose that differences in these parameters may serve to highlight potential epigenetically mediated changes in regulatory tuning separating phenotypes. After conducting a Wilcoxon-signed rank test on update speed and a chi-square test on detection thresholds, we found the vast majority of these to be statistically different (Bonferroni-corrected p<<0.01) between the two model populations. Indeed, 129 out of 133 update speeds and 671 out of 676 detection thresholds showing statistically significant differences across phenotypes. This result is not all that surprising given the large number of models in each pool and the correspondingly high degrees of freedom supporting a high significance to even small changes. To focus more specifically on the most meaningful variations in regulatory programming separating phenotypes, we also examined the magnitude of differences in update speed and threshold, both independently and in combination.

To draw out the largest differences in update speed separating the LSev and MSev model populations, we isolated nodes where differences in average update speeds across phenotypes were equal to or exceeded the mid-range threshold of 6 units (δ≥6) (**Fig. 3A**). The highest divergence in average update speed corresponded to nodes mitogen-activated protein kinase activated protein kinases (MAPKAPK5), 5-azacytidine-induced protein 2 (AZI2), CCAAT/enhancer-binding protein Δ (CEBPD), C-C motif chemokine ligand 5 (CCL5), epiregulin (EREG), and Forkhead box protein (FOX)O4 (δ > 9). Nodes organize into sets sharing a common differential in update speed with the size of these node sets increasing the magnitude of this difference decreases (**Fig. 3B**). Ultimately, we find 4 nodes exhibiting virtually identical update speeds with differences failing to achieve significance, namely MAPK10, IL-6, Nucleotide-binding oligomerization domain-containing protein 2 (NOD2), and IL-18 (δ < 0.01). A similar assessment can also be applied to network relationships that transmit regulatory actions from one node to another when activation of the upstream mediator exceeds a detection threshold. This discrete value parameter or categorical parameter is set to 1 in the case of a more potent mediator and more sensitive downstream target and 2 in the opposite case of a weaker mediator and less responsive target. Phenotypic differences in edge detection threshold were found by tallying the number of high and low detection thresholds for each relation and assessing overrepresentation of more permissive versus more refractory signal transmission in each model pool (**Fig. 4A**). Though not strictly unanimous, opposing settings of low versus high threshold were shared by over 97% of the models in each pool for 276 out of 676 total relationships. Among these 276 differentially permissive relationships, 141 were assigned a high threshold and the remaining 135 a low threshold value in the model pool describing the LSev phenotype. Conversely, a similar near unanimous agreement across models in each pool supported identical detection threshold settings for 311 relationships (**Fig. 4B**) or 46% of all regulatory relationships. Once again as with update speed, we find that while differences exist, regulatory similarities outnumber the latter and that the dramatically different outcomes manifested by each phenotype may arise from dysregulation in a relatively focused set of processes.

**Figure 3:**
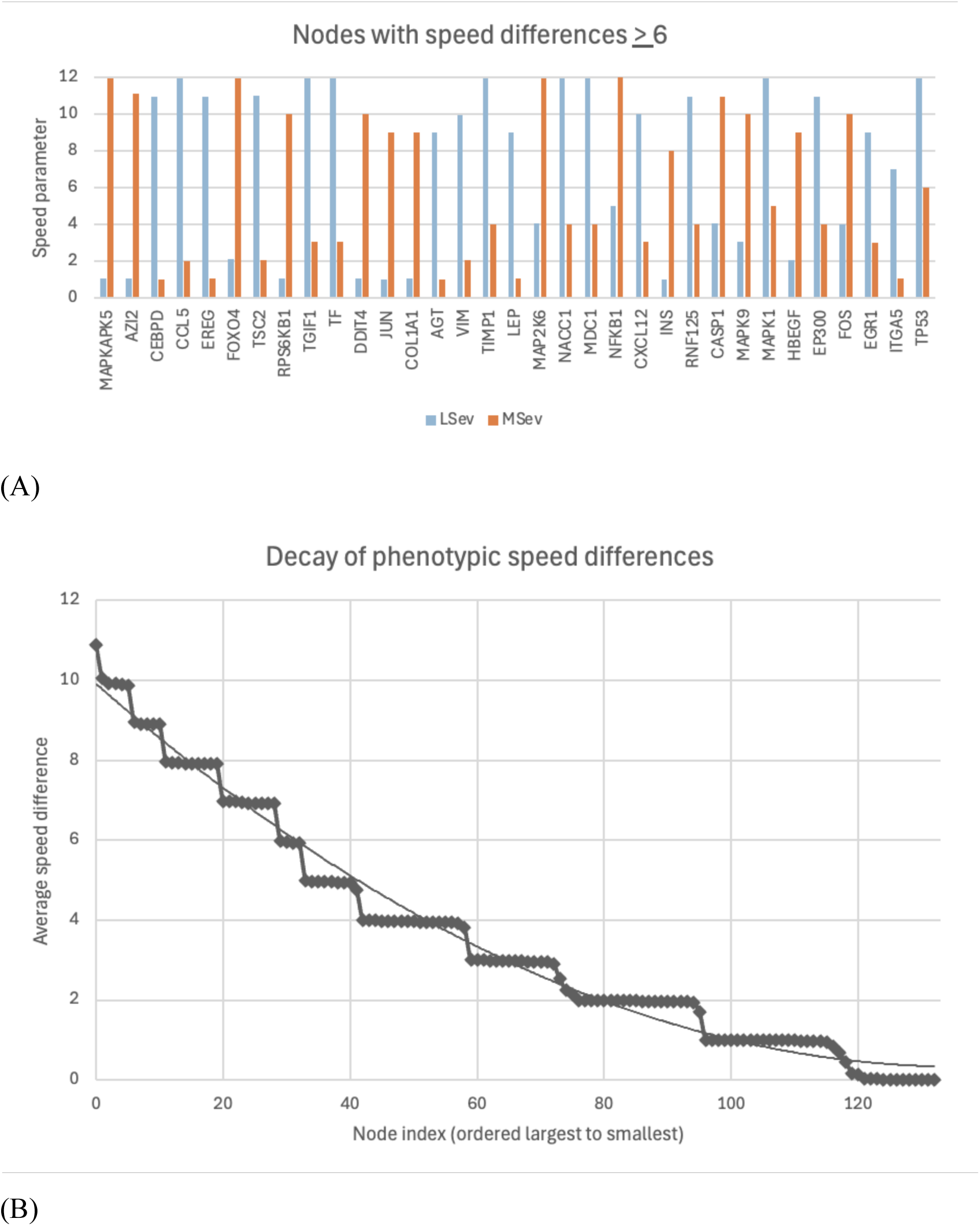
Phenotypic variation in nodal speed. A) Differences between population averages of nodal speed are plotted here (shown are only those with difference >6). B) The decay pattern of update speed differences. The broadening of the bottom three quarters of the staircase highlights that an increasing number of nodes are recruited into groups sharing smaller differences in update speed and vice versa.

**Figure. 4:**
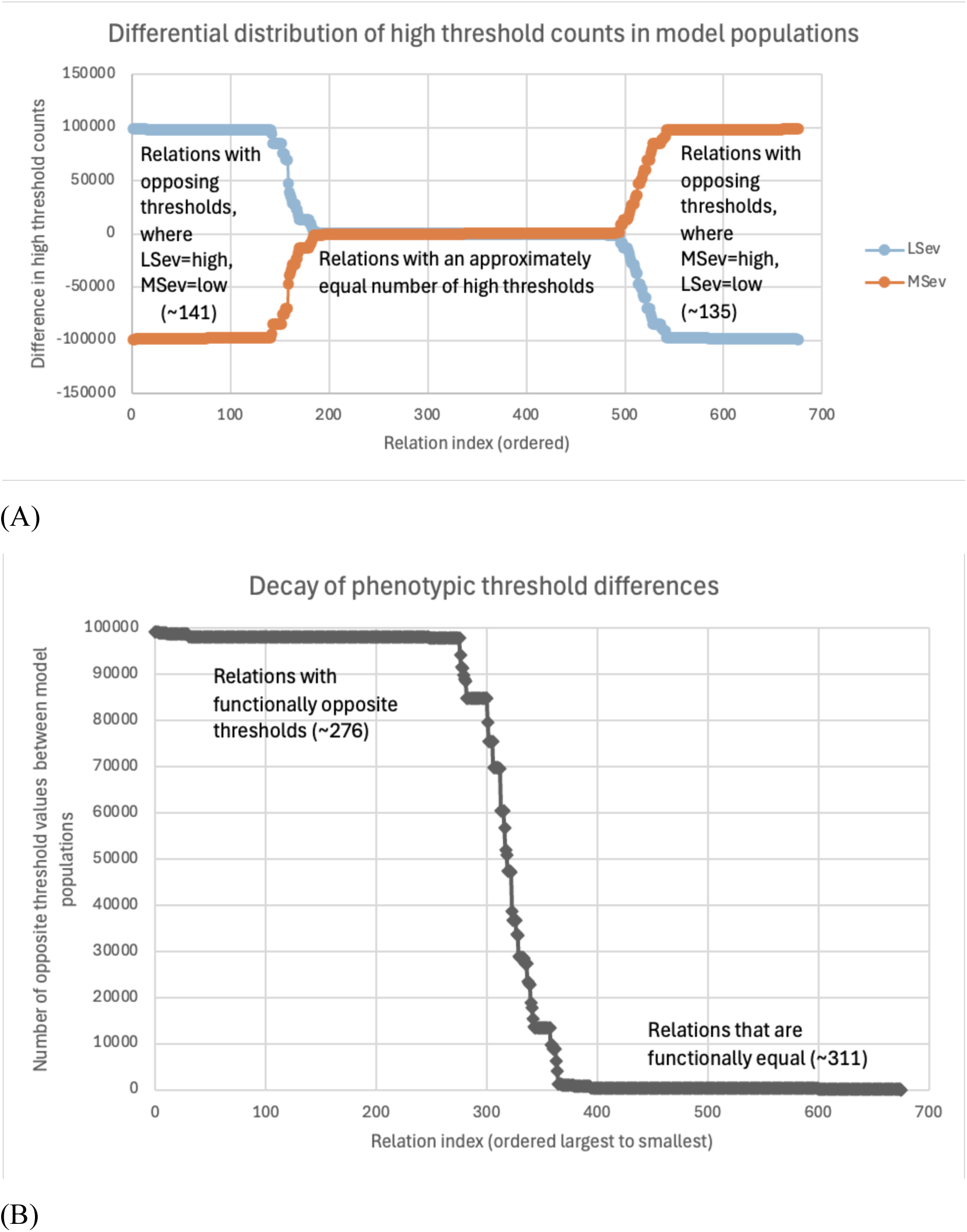
Phenotypic variation in thresholds. A) Differences in high threshold counts are plotted here, relations ordered from greatest to least difference (from LSev perspective). B) The decay pattern of threshold differences, ordered from largest to smallest in absolute difference.

Both update speed and detection threshold will influence how frequently nodes may update their activation state. If a node is assigned a high update speed, then it is more likely to change state frequently, assuming signals from upstream mediators are detected at a low threshold and drive a new target state. Depending on their settings, these two parameters can act in unison to promote rapid dynamics or in some cases counteract one another. For example, a node may be frequently available for change (i.e. high update speed) but not receive a new target state very frequently because it is relatively insensitive to incoming regulatory prompts (i.e. high detection threshold for incoming signals). Conversely, opposite settings of low upstream detection thresholds and high update speeds combine cooperatively to deliver fast kinetics in that protein node (and vice versa). To highlight those nodes for which phenotypic differences in update speed and detection threshold work cooperatively, we derived a combinatorial scoring scheme along two axes, namely slow-fast and protective-maladaptive, to describe their combined effects. Here, a slow score is associated with low speeds and high upstream thresholds, whereas a fast score denotes high speeds and low thresholds. A protective score describes strong representation in the LSev phenotype pool, whereas a strong representation in the MSev model pool is considered maladaptive. As a large proportion of commonly studied epigenetic changes result in muted signaling, we focus here on slow-protective (SP) scores and slow-maladaptive (SM) scores. To assign SP and SM scores, for each node we multiplied the number of upstream phenotypic differences in detection threshold by its phenotypic change in update speed. Nodes where regulatory differences in speed and threshold act cooperatively will be associated with large negative scores. (see Methods). Based on these scores, protein nodes where slowed or muted regulation was protective and led to a slower progression ((i.e., large negative SP score, <-20) included IL-1β, TNFα, and FOXO4 (**Fig. 5A**). Accordingly, these same nodes are associated with a weak maladaptive response (large positive SM score, >20). Visibly less distinct than the top SP nodes are the top SM nodes, prostaglandin-endoperoxide synthase 2 (PTGS2) which codes for cyclooxyegenase 2 (COX-2), and vimentin (VIM) (**Fig. 5B**). Nonetheless, as these are also associated with moderately weak SP scores of 11.9 and 7.9 respectively, a more muted regulation of these proteins may at least in part contribute to a poorer outcome.

**Figure 5:**
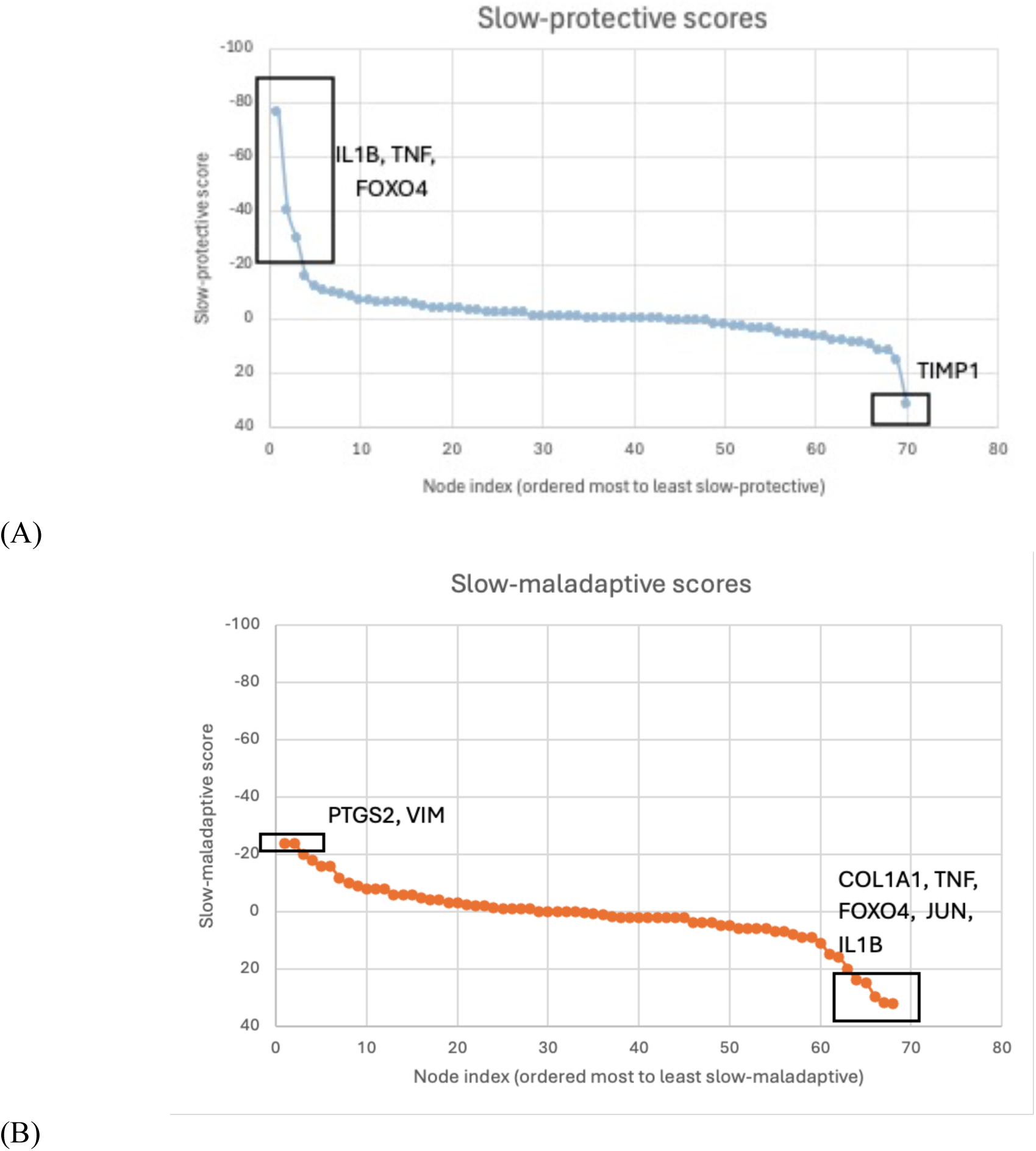
Slow-protective and slow-maladaptive scores reveal nodes with regulatory variation. A) Distribution of slow-protective scores among nodes that have opposing upstream thresholds where LSev is high and MSev is low. In this case, a large negative score indicates strong association between regulatory dampening and better experimental outcome. IL-1β, TNFα, and FOXO4 stand out with substantially higher scores than other nodes. B) Distribution of slow-maladaptive scores among nodes with opposing upstream thresholds where MSev is low and LSev is high. A large negative score indicates association between regulatory dampening and a worse experimental outcome. Although less distinctive than the top slow-protective nodes, this plot shows PTGS2 and VIM to be the most maladaptive when slowed. (Note the reversed y-axis in both).

**Figure 6:**
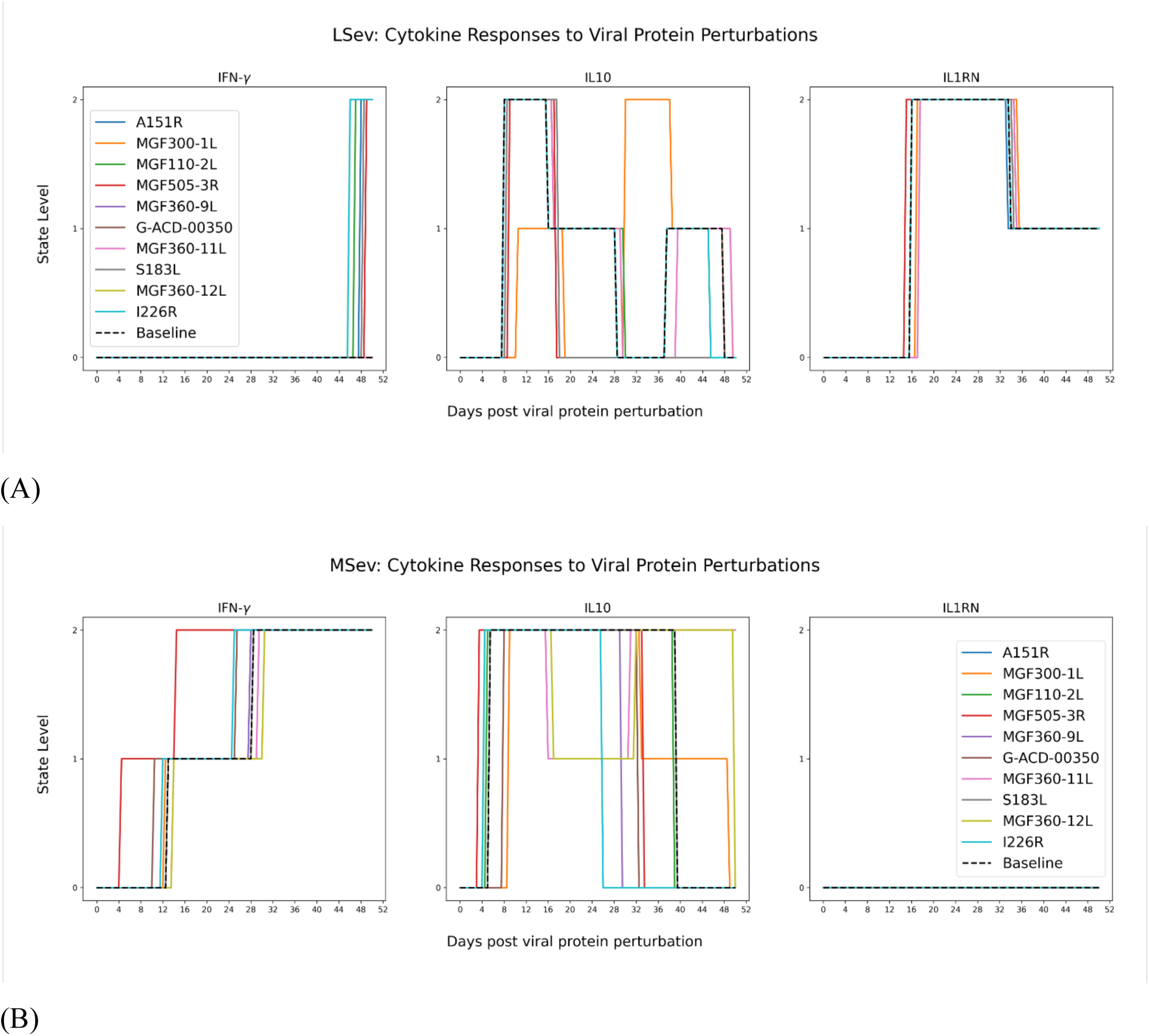
Simulated response to viral protein challenge. Simulated response in IFN-γ, IL-10 and IL-1RN to challenge where each of the 10 individual viral proteins selected here as strong interactors are introduced at t=0 and held at the maximal activation level of 2 until t=4 time steps or 2 days then released.

#### Functional impacts on immune response

In an attempt to explore the functional repercussions on immune response of these differences in regulatory tuning, we conducted simulated perturbation experiments where we systematically challenge the regulatory network in each phenotype with individual pathogen proteins. For each challenge, one of the 10 viral proteins retained here as strong interactors are introduced at the maximum activation level of 2 and held constant at that level (overabundance conditions) for t=0 to t=4 time steps of 12 hours (2 days), then released for the remaining time steps. Though survival of the study animals did not exceed 8 days, we simulated an extended immune response of up to 52 days as an additional means of verifying model stability. These challenges are applied to what may be regarded as the single most representative model for each phenotype, or the model closest to the centroid of the parameter space for that model pool. In the case of the LSev phenotype, this model is separated from the centroid of the parameter space by a Hamming distance of ∼25 bits (88.16 Manhattan), while its counterpart for the MSev phenotype sits at a Hamming distance of ∼7 bits (39.36 Manhattan) away from that pool’s centroid. As expected, the simulated challenges confirm in both phenotypes a muted and delayed IFN-γ response to the subset of 10 viral proteins selected here. Indeed, the earliest IFN-γ response in the LSev phenotype is predicted to occur at 45 days post-challenge (I226R) or later, far exceeding the 8-day survival time. In the case of the MSev phenotype however, viral protein multigene family (MGF)505-3R produces a predicted IFN-γ response at day 4, or 2 days prior to death (day 6). This response is followed by model-predicted IFN-γ responses to ASFV genome G-ACD-00350 and the remaining viral proteins after day 10.

Turning our attention to immune suppressive actions of these proteins, we find a predicted increase in IL10 expression occurring at day 7, with increased expression of IL-1 receptor antagonist N (IL1RN) predicted at day 14 in the LSev phenotype, or long after death. This IL1RN response in LSev is shared almost identically across all 10 viral proteins including MGF505-3R. Once again, in the case of the MSev phenotype we find an earlier response to challenge with IL10 activation by MGF505-3R being recruited at day ∼3, followed shortly thereafter by predicted IL10 responses to viral proteins I226R and MGF110-2L. For the MSev phenotype this IL10 response does not coincide with a recruitment of IL1RN with the latter remaining at basal levels across the simulation horizon of 52 days.

## Discussion

The reality of working with high containment level pathogens in living animals is that data are logistically challenging to acquire. Study populations tend to be low, with the number of samples that can be taken and observable markers available for analysis also being limited [21-23]. However, we have shown here that a great deal of information can still be extrapolated from small amounts of data when a suitable modeling framework is applied. In recent work, we have shown that by leveraging our prior knowledge of network biology and aligning this with whatever limited data might be available as examples of expected behaviors, we can bring out key regulatory pathways and mediators that offer a mechanistic basis for explaining two seemingly different phenotypes in persistent respiratory illness[24]. Here we apply the latest extensions to this approach to explore underlying immune regulatory mechanisms that might explain differences between two distinct courses of illness progression in ASFV infection. Using a published pathogen-host protein interactome [17] together with extensive text and schema mining we assemble a closed loop regulatory feedback network. As these are genetically standardized pigs of the same species [21], we expect that this fundamental immune signaling network structure will be conserved and that divergent illness trajectories emerge as a result of epigenetic adaptations that will be reflected by a phenotype specific tuning of the logic and timing of immune decisions. With progression in each phenotype being only sparsely sampled and very partially observed, we find large populations of competing regulatory programs in each phenotype capable of directing the flow and timing of immune signaling such that they support dynamic behaviors that include those observed experimentally. Interestingly, the correspondingly large sets of competing network share very strong commonalities with over 90% of structural and kinetic settings being preserved across the near totality of the model sets for each phenotype. This would suggest that differences in illness course may arise for relatively focused changes in immune signaling with these large sets offering high statistical resolution in highlighting details in immuno-protective/maladaptive regulatory variation. Although the logic modeling framework used here does not explicitly represent epigenetic modification of regulatory tone, we propose that model parameters such as detection threshold and update speed may be considered viable analogs. Scoring the combined effects of changes in these model parameters, we find important alterations in the regulation of IL-1β, TNFα, and FOXO4 across phenotypes with dampening of these responses being characteristic of a slower illness progression. In the cases of IL-1β and TNFα, about two thirds of these altered upstream regulators are activators, and one third are inhibitors. For FOXO4, we find an opposite ratio with two altered regulators being inhibitors and one being an activator.

Consistent with our analysis, elevated IL-1β has been associated with increased severity in viral infections [25] with IL-1β inhibitors emerging as a potential treatment component in severe COVID-19 [26,27]. Additionally, the nuclear factor κβ (NF-κβ) and NOD-like receptor pyrin domain-containing protein 3 (NLRP3) inflammasome pathways that promote IL1β release are known to be highly susceptible to DNA methylation supporting chronic inflammation [28,29]. This susceptibility of upstream IL-1β activators to epigenetic modulation further supports the hypothesis that this may be a key mechanism in accelerating illness progression in ASFV infection. Likewise, our models also suggest slowed TNFα activation may led to a slower progression. A strong promoter of cell death, TNFα recruits the simultaneous activation of the NF-κβ pathway, leading to further cytokine release and inflammation [30,31], often ending in cytokine storm. Indeed, the administration of TNF neutralizing antibodies to Severe Acute Respiratory Syndrome Coronavirus 2 (SARS-CoV-2) mouse models resulted in a more controlled cytokine response and a reduced mortality rate [32]. Furthermore, there is evidence that epigenetic mechanisms including histone methylation and acetylation are implicated in regulating TNFα expression, and that these occur both developmentally and acutely in response to stimuli [33]. As with IL-1β, environmentally and developmentally induced adaptations in TNFα regulation may be an impactful contributor to progression. In contrast to IL-1β and TNFα, the majority of alterations to FOXO4 regulation consisted of a more muted inhibition, implying that the presence of FOXO4 may somehow be protective. Interestingly, this has been observed in the contexts of other viral infections, especially DNA viruses. FOXO4 has been reported to antagonize the transcription and replication of the DNA virus Hepatitis B [34] and the retrovirus Human immunodeficiency virus 1 (HIV-1) [35]. Furthermore, FOXO4 is a powerful antioxidant with FOXO4-knockout in a mouse model of inflammatory bowel disease resulting significantly higher levels of inflammatory cytokines including TNFα and IL-1β, due at least in part to the loss of FOXO4’s antagonism of NF-κβ induction [36]. Classic epigenetic mechanisms regulating FOXO4 remain poorly understood, however, there are several instances of post-transcriptional regulation of FOXO4 via microRNAs, non-coding RNAs that are themselves epigenetically regulated and implicated in epigenetic feedback loops [37-39].

Interestingly, and perhaps not surprisingly, these regulatory differences affecting illness progression were observed in simulations to also result in altered responses to challenge with individual viral proteins. Certainly, the pursuit of subunit vaccines for ASF continues to be a challenging endeavor plagued so far by limited effectiveness and inconsistent results [40]. Here we attempt to use simulated challenges as a potential means of rapid screening for potential immunogenic proteins of interest. Among the limited set of viral proteins considered in this proof-of-concept work, MGF505-3R was predicted have a immunostimulatory effect on IFNγ response but only in the case of the more severe progression phenotype (MSev). Likely due to the high virulence of this gene group, MGF505-3R, has not been tested in a subunit vaccine to our knowledge. However, this contribution to high virulence has made it a target of interest for deletion-based attenuated vaccines [41,42]. Although in a recent study, MGF505-3R was associated with inhibition of IFN-γ response [43], this was based on an acute early response (24 hours vs. 4 days) in a HeLA cell assay rather than *in vivo* swine. Other viral proteins in our model that have shown immunogenicity in previous studies include MGF360-11L [44] and A151R as part of an adenovirus-delivered antigen cocktail [45]. Though neither of these produced noticeable effects on IFN-γ in either phenotype’s model when introduced individually, our continuing work will include the study of synergistic effects induced by combining multiple viral protein antigens. Such future modeling work will also necessarily address the co-selection and introduction of adjuvant immunostimulants, with the latter requiring the inclusion of added granularity in first-responder proteins such as Toll-like receptors (TLRs) and a broader set of candidate ASFV proteins.

While any computationally generated hypothesis should be verified with the appropriate experimentation, this study demonstrates how prior knowledge and principles of logical reasoning can be leveraged to infer broad regulatory dynamics of biological networks from experimental data where physical samples are scarce, difficult to collect and assess. As a proof-of-concept, we provide putative evidence supporting the hypothesis that differences in ASF progression may arise from adaptive epigenetic changes, particularly in the regulation of IL--1β, TNFα, and FOXO4. Insight gained in this way through computer simulations may serve to develop focused experiments that best advance our knowledge base, informing on measures for ASF control including vaccine design. Moreover, the ability to mimic these changes and the corresponding variability on immunogenic response to viral proteins may offer a novel opportunity for conducting rapid and large-scale screening for adjuvant-antigen co-selection, with an added focus on compounds that can modulate such epigenetic changes [46].

## Methods

### Construction of the regulatory network

The ASFV host-pathogen interaction network used in this study was derived from the protein-protein interactome published by Wu et al. [17], consisting of 590 host proteins, 77 viral proteins, and 8946 undirected interactions. To reduce this number to a manageable amount and tease out the most significant contributors, Otsu’s method [47], a thresholding algorithm, was applied, leaving 35 proteins and 34 interactions. Using these 35 proteins as the starting point, a number of their first neighbors was recovered from the BEL Selventa Large Corpus [48] and the BioPax/Pathway Commons [49] and added back into the network. After this step, the network was comprised of 279 proteins from both host and ASFV, and 931 relationships. At this stage, some pruning was done to ensure the network was a true regulatory network in that it was a closed loop system and therefore fully self-regulating (no sources or sinks). To achieve this, every node in the network needs to have at least one input and at least one output. After removing the proteins that did not satisfy this criterion, the size of the network was reduced to 133 proteins and 588 relationships.

A dynamically responsive regulatory network requires that all proteins must have the capacity to be both positively and negatively regulated. The modeling framework has an inherent bias term of 0 at every node, meaning each node has the chance to be negatively regulated if there are no positive inputs to it. However, positive regulation can only occur by upstream mediators; therefore, each node must have at least one upstream positive regulator. To find additional positive regulators for the 23 proteins that were so far being exclusively downregulated, manually curated text mining of the EmBiology database was carried out, which resolved 77 upregulating relationships for 20 of the 23. For the remaining 3, the CleanLab Trustworthy Language Model was used to predict positive regulators. The CleanLab TLM is a generative AI model that provides a trustworthiness score for the answers it provides [50]. Out of the 11 predicted relationships that were added to the network, 2 had trustworthiness scores of 80%, and the others had scores of 90%. At this point, the network presented a fully closed-loop architecture where every node could also be positively and negatively regulated. The final resulting network shown in **Fig. 1** has 133 proteins, 10 of which are viral, and 676 relations.

### Generation of immune response programs

#### Modeling framework

The modeling framework employed in this study is the discrete logical formalism of regulatory networks. While applied here to ASFV, it is largely problem-agnostic and has been described in detail in previous works [51-54]. Briefly, biological regulatory networks are represented as a collection of “nodes” (biological mediators, e.g., proteins, signaling molecules, etc.) and “edges” (the interactions between mediators). State values of nodes can take on one of three discrete values at a given time step:

- *s*_*i*_ ∈ {0, 1, 2} is the state value of entity *i*, where 0 is associated with low, 1 with moderate, and 2 with high levels of that entity.

Each interaction (edge) has three parameters associated with it. Where the source node is *i* and target node is *j,*

- *w*_*ij*_ ∈ {0, 1, 2} is the weight of edge *ij*, where 0, 1, and 2 denote non-existent, moderate or strong interactions respectively.
- *p*_*ij*_ ∈ {−1, 0, 1} is the polarity of the edge *ij*, where -1 denotes an inhibitory effect, 1 an activating effect, and 0 a non-existent one.
- *t*_*ij*_ ∈ {1, 2} is the threshold that denotes the minimum level of *s*_*i*_required for the interaction *ij* to be active.

Together, these parameters contribute to determining each node’s target state, or “image”, at any given time step, wherein the image *K_j_* of some node *j* is the cumulative input of all source nodes *i* ∈ *I_j_* that have *j* as a target:

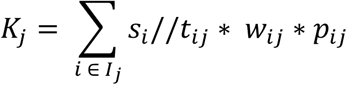

Each node that is not already equal to its image is placed in a priority queue and is updated according to the method outlined in Lyman et al. [55].

This requires the definition of three additional nodal kinetic parameters:

- *m*_*i*_ ∈ {0, 1, 2, 3} is the memory parameter of *i*, i.e., how many consecutive time steps the node *i* must be told to update before transitioning to the prescribed state.
- *u*_*i*_ ∈ [1, 12] is the speed of action of node *i*. These parameters increase or decrease a node’s priority to be updated.
- *d*_*i*_ ∈ {1, 2} is the step size, the maximum number of state levels *i* can change per time step.

If a node is eligible to update, meaning its image is different from its current state and its memory criteria has been met (*m*_*i*_), then its priority is computed as the product of its speed *u*_*i*_ and how many time steps it has been since it was last updated. The nodes with the highest priority value are updated at the end of each time step, moving a maximum of *d*_*i*_ steps toward their image. Within this modeling framework, a complete model is given by the vector, **x**, where:

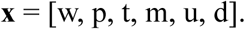

#### Encoding of experimental data

The experimental data used as reference for model generation is acquired from a study by Zuo et al. [21]. In the study, six pigs are infected with ASFV, and several biomarkers are sampled over time before the pigs die by cytokine storm or are culled at the humane end point. The study distinguishes between two observed phenotypes—one group of pigs that died after 6 days post-infection, and another that died after 8. In this study, we refer to the former case as “More Severe” (MSev) and the latter as “Less Severe” (LSev). We identify two separate families of models that capture each phenotype’s distinct cytokine trajectories. The cytokines whose trajectories are used are those in common with our constructed closed-loop network: IL-1β, IL-6, TNFα, IL-4, and IL-10. The absolute cytokine measurements reported in the paper are normalized within each trajectory and binned into three categories (0, 1 or 2, i.e., Low, Medium or High), allowing for the experimental data to be interpreted by our modeling framework. Another piece of experimental data from the paper that is used to guide model generation is blood viral nucleic acid volume. While this cannot be a hard constraint because specific viral proteins are not measured, the objective function is designed to reward models that exhibit an average viral protein load similar to what was seen experimentally; that is, “Low” for the first 3 days, “Medium” at Day 4, and “High” by Day 6.

#### Determination of initial conditions and time step

The model vector **x** offers the ability to simulate the trajectory of some condition over time. To implement this practically, initial conditions and time step sizes are also required to be specified. In our case, we must set initial conditions for all nodes in the ASFV network, despite known baseline values only being available for the five measured nodes. Rather than randomizing, we take a more systematic approach of formulating a constraint satisfaction problem to determine each phenotype’s pre-infection steady state. Utilizing the CPMPy constraint solver library [56], the closed-loop regulatory network logic and modeling framework described above are input as constraints. Additional constraints require that the five measured nodes take on the pre-infection values reported in Zuo et al., and the viral proteins be at “0”. The solutions to this constraint satisfaction problem will include possible steady states for all network nodes and the corresponding model parameters that make that state possible. The steady states from the first 100 solutions are taken as the initial condition for the simulations, with the corresponding steady state models being used to initialize the optimizer with structurally valid starting guesses. The optimal time step size is calibrated through sampling. Abridged optimizations with step sizes of 2 days, 1 day, 12 hours, and 6 hours were carried out. A step size of two days is the same as the time increment of the data, but for step sizes smaller than this, the intermediate states are free variables and allowed to take on arbitrary values. The quality of solution plateaued at a step size of 12 hours, with a further decrease in step size not resulting in further improvements; therefore, a step size of 12 hours was used for the subsequent full-scale optimizations. This means that 12 time steps are required to simulate the 6-day MSev trajectory, and 16 for the 8-day LSev trajectory.

#### Global search for immune regulatory parameter sets

In this study, we aim to find at least one parameter set **x** that results in simulated data that closely matches the transcribed reference data. With bounds on each parameter as outlined above (“Modeling Framework”), the network’s 676 edges and 133 nodes give rise to a vector size of 2,427 parameters. There are 1.6x10^1112^ possible combinations of these parameter values; however, because 579 polarities are previously known, this number is reduced to 9 x 10^835^. Due to the large search space size and discrete nature of the modeling framework, we require a robust global optimization method. For this, we employ the simulated annealing algorithm, implemented in parallel for GPU architecture. This algorithm is used widely for many global optimization problems [57-59]. Its success is largely due to its ability to balance exploration with exploitation; that is, it is willing to temporarily accept a sub-optimal answer in order to avoid being trapped indefinitely in a local minimum. The probability with which the algorithm accepts a worse answer follows a cooling curve. In our optimizations, all computations use a starting temperature of *T* = 1000 and a cooling factor of 0.99. During a temperature cycle, each GPU thread performs 5000 search iterations and sends an optimal solution back to the host CPU. The best-performing solution out of all threads is then broadcasted to the other threads, and the next cycle begins. The optimization is stopped when the temperature reaches 0.001. A more technical description of the optimization algorithm and benchmarking can be found in a previous work [20].

#### Optimality of regulatory parameter settings

The quality of a solution is determined by its error; that is, by the distance computed in the Manhattan metric between the simulated data it produces and the transcribed reference data. The quality is further determined by whether it produces an average viral protein trajectory that is in line with the blood viral nucleic acid volumes presented in the reference study [21]. If the average is approaching the target level, the sum total of viral nucleic acids detected across of all viral proteins is subtracted from the solution’s error, and if it is at the target level, the sum is doubled before subtracting. This has the effect of guiding the optimization algorithm to favor solutions with non-zero viral protein trajectories, even though other solutions with little to no viral activity may still technically capture the reference data with little or no error. This results in the following objective function, which we seek to minimize. Let *t* ∈ {0, …, *T*} denote each time step of the simulation and *j* ∈ {1, … *N*} denote the index of each node. We then define *s_t,j_*(**x**) to be the simulated state of node *j* at time *t* given some candidate parameter vector **x**, and *r_t,j_* to be the corresponding reference data for node *j* at time *t*. The absolute difference between these values, summed over all states for which *r_t,j_* exists, is the error term, Δ_t_, at time *t*. The second term, Θ, is a regularization term dependent on the time step and the average viral state at that time step, *V̄*_*t*_ . If the time step is one where an ideal viral load has been prescribed (i.e., Days 4, 6, and 8; see above) and the average viral state is either approaching or equal to the prescribed ideal viral load, then *ρ* is non-zero (*ρ =* 1 if approaching, *ρ* = 2 if equal). The weight *ρ* is amplified by the sum of the *N_v_* viral protein states at time *t*. These two terms are summed over all time steps and subtracted:

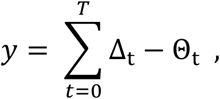

where

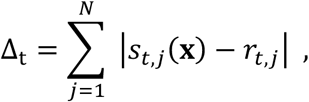

and

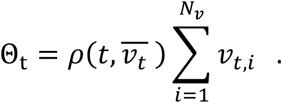

### Analysis of resulting models

With such a large search space and sparse data to fit, it is likely that once the optimization converges to some minimum there will be far more than one solution that results in the minimum value. Indeed, in this case, we find many models all resulting in the same objective function value for each phenotype’s optimization. Despite the large number of solutions, we can still draw conclusions about phenotypic differences by analyzing differences between each phenotype’s resulting solution set.

#### Statistical analysis of kinetic parameters

To perform the population comparison, 100,000 models that give the minimum error are collected for each phenotype (Δ = 1 for LSev, Δ = 2 for MSev). The parameters of interest are the speeds and thresholds; these provide information about regulatory kinetics, pointing to sites of differential regulatory amplification or dampening. These parameters are first checked for statistical difference between the two model populations. Given the binary nature of the thresholds, a chi-square test is carried out. With an original α= 0.01 and a Bonferroni correction resulting in α =1.48e-05, it is determined that 671 out of the 676 threshold parameters are statistically different. A Wilcoxon signed-rank test is used for the speeds, given the wider range but still ordinal data, and this shows 129 out of 133 speeds are statistically different (corrected α= 7.52e-05).

#### Defining slow-fast/protective-maladaptive scores

Because there are statistically significant differences between almost the entire set of kinetics parameters of the two phenotypes, further processing of both parameters must be done to draw out nodes that are the most differentially regulated. Speed and threshold are looked at independently, and then in combination. For the former, the average speed of each node within each phenotype is determined, and nodal differences between the two phenotypes are calculated. For phenotypic differences in threshold, the ratios of high vs. low thresholds are computed for each relation. The thresholds with the largest phenotypic differences (i.e. the relations where a proportion >97% of each phenotype’s models point to opposite thresholds) are considered functionally distinct and are treated as inequalities in the subsequent scoring method. Because the speed of a given node and the thresholds of the upstream regulators acting on that node are antagonistically related (e.g., a high nodal speed and high upstream regulatory thresholds effectively work against each other), we derive a scoring scheme along two axes—slow-fast and protective-maladaptive—to quantify the combined effects of speed and threshold on experimental outcome. Here, kinetics parameters found primarily in the LSev models are considered “protective”, and those found in MSev models are considered “maladaptive”. These axes can be combined in four ways, giving four scores: slow-protective (SP), fast-maladaptive (FM), slow-maladaptive (SM), and fast-protective (FP). The first two are equivalent, as are the last two (why?), and so we choose to limit our perspective to the effects of slowed regulation. This leaves the slow-protective (SP) and slow-maladaptive (SM) scores to be computed.

For each node, *k,* where *i* is an upstream regulator of *k, n_k_* is the total number of *k*’s regulators, *t_i,k_* is the threshold value of *i* acting on *k* (given >97% of models agree), and *u_k_* is the average speed of *k*, these scores are defined as

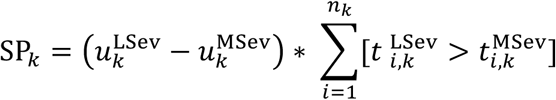

and

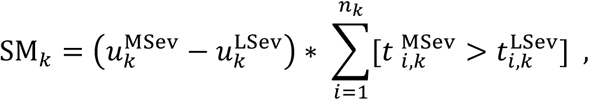

with the notation

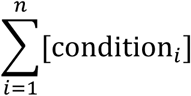

referring to a count of how many times condition *i* is found to be true. In both cases, a large negative speed differential indicates phenotype-dependent slower activity for node *k.* This value gets amplified by the second term when there is a greater number of upstream regulators acting to further enforce slower activity (i.e. phenotype-specific higher thresholds). Therefore, a *large negative* SP score indicates a node that is protective when slowed (or, equivalently, that is maladaptive when quickened), and a *large negative* SM score indicates a node that is maladaptive when slowed (or, equivalently, that is protective when quickened).

#### Software and hardware requirements

The optimization code is implemented in Python 3.12. Libraries used include NumPy [60], pandas [61,62], CPMPy [56] to initialize the optimizer, Numba [63] to compile the code and interface with CUDA [64] and the GPU device, and SciPy [65] for statistical analyses. All model assembly, parameter identification and numerical experiments were conducted on compute nodes provided by the Centre for Quantum Topology and Its Applications, as part of the Copernicus cluster supported by the University of Saskatchewan’s Advanced Research Computing (ARC) team. Each compute node consists of 32 CPU cores (Intel Xeon Platinum 8356H CPU @ 3.90GHz) with 1.5TB of RAM and 20TB of highspeed NVMe local storage housed with 4 Nvidia A100 GPU with 40GB VRAM. All CUDA kernels are submitted with 10,240 threads in block sizes of 256.

## Supporting information

In file **Supplementary_tables.xlsx** see tabs:

**S1 Table**. Regulatory relationships defining ASFV pathogen-host network. (XLSX)

**S2 Table**. Pre-infection steady state constraints less severe (LSev) and more severe (MSev) phenotypes. (XLSX)

**S3 Table**. Less severe (LSev) phenotype cytokine trajectory during ASF infection. (XLSX)

**S4 Table**. More severe (MSev) phenotype cytokine trajectory during ASF infection. (XLSX)

**S5 Table**. Phenotypic differences in average node update speed. (XLSX)

**S6 Table**. Decay of differential in average node update speed across phenotypes. (XLSX)

**S7 Table**. Phenotypic differences in regulatory relationship (edges) detection thresholds. (XLSX)

**S8 Table**. Decay of threshold differences across phenotypes. (XLSX)

**S9 Table**. Slow-protective scores. (XLSX)

**S10 Table**. Slow-maladaptive scores. (XLSX).

## Author Contributions

**Conceptualization:** Joyce Reimer, Sureesh Tikoo, Gordon Broderick

**Data Curation:** Joyce Reimer, Pranta Saha, Kira Comfort

**Formal Analysis:** Joyce Reimer, Pranta Saha, Kira Comfort

**Funding Acquisition:** Heather Wilson, Gordon Broderick

**Investigation:** Joyce Reimer, Pranta Saha, Kira Comfort

**Methodology:** Joyce Reimer, Pranta Saha, Gordon Broderick

**Project Administration:** Sureesh Tikoo, Heather Wilson, Gordon Broderick

**Resources:** Connor Burbridge, Brook Byrns, Steven Rayan, Gordon Broderick

**Software:** Joyce Reimer, Pranta Saha, Gordon Broderick

**Supervision:** Gordon Broderick

**Visualization:** Joyce Reimer, Gordon Broderick

**Writing – Original Draft Preparation:** Joyce Reimer, Gordon Broderick

**Writing – Review & Editing:** All authors

## Acknowledgements and Funding Information

This work was supported by the University of Saskatchewan’s Centre for Quantum Topology and Its Applications (quanTA) and by the Vaccine and Infectious Disease Organization (VIDO). VIDO receives operational funding from the Canada Foundation for Innovation (CFI) through the Major Science Initiatives Fund and from the Government of Saskatchewan through Innovation Saskatchewan and the Ministry of Agriculture. The quanTA Centre’s high-performance computational work has been advanced through a CFI John R. Evans Leaders Fund grant while access to quantum computing resources through IBM Quantum and PINQ2 has been made possible by a PrairiesCan Regional Innovation Ecosystems (RIE) contract (both awarded to SR). H.L.W. is supported by a Natural Sciences and Engineering Research Council of Canada (NSERC) Discovery Grant (416587). We thank the University of Saskatchewan Advanced Research Computing (ARC) team for their efforts and responsiveness in creating an excellent local environment for this computational work. This article is submitted with the permission of the Director of VIDO.

